# The intestinal immune response is influenced by nutritional-status and increased physical activity level

**DOI:** 10.64898/2026.04.01.715829

**Authors:** Cecilie Bæch-Laursen, Nora Silvana Nägele, Beckey Trihn, Carolina Manzano, Sara Vargas, Anna Heberg Johnson, Helga Ellingsgaard, Joel Vizueta, Benjamin Anderschou Holbech Jensen, Bente Klarlund Pedersen, Paula Sanchis

## Abstract

Beyond its role in digestion and barrier function, the intestine is an energy-responsive organ that actively regulates molecular metabolism. Whether and how lifestyle interventions regulate intestinal metabolism at both tissue and molecular levels remain unclear. Here, we show that both chronic exercise and dietary energy density drive robust, segment-specific intestinal remodeling. Voluntary wheel running in *ad-libitum* chow fed mice, induced elongation of the small intestine and colon, alongside pronounced, region-specific, transcriptional changes in the proximal, mid, and distal small intestine, particularly within immune and stress-related pathways. Caloric dilution diet also increased intestinal length in mice but elicited transcriptional adaptations, prominently in the proximal small intestine, directly linking energy density and intake to structural and molecular plasticity. In contrast, voluntary wheel running in control-fed and caloric-diluted-fed mice subtly modulated immune-associated gene expression, highlighting that diet and physical activity induce complementary and mechanistically distinct effects on the gut. We further identified an exercise-induced state of intestinal preconditioning. Upon refeeding, sedentary mice mounted robust, segment-specific activation of apoptotic, proliferative, and immune pathways. Similarly, acute treadmill exercise acted as a transient intestinal stressor in sedentary animals by shortening the length of the small intestine and rapidly activating epithelial stress, apoptosis, proliferation, and immune signaling. However, these responses were attenuated in chronically active mice despite higher basal expression of key genes, consistent with adaptive epithelial remodeling. The results suggest that habitual physical activity buffers acute nutritional stress and restrains excessive intestinal immune activation. Finally, translational plasma analyses in humans demonstrate that acute moderate-intensity exercise increases circulating markers of monocyte activation and epithelial stress, including CD14, IL-32, Reg-3-alpha and I-FABP, in both lean and obese individuals. Collectively, these findings suggest that the intestine plays a role as a metabolic organ that integrates energy-sensing signals from diet composition and physical activity.

## 1. Introduction

Research of the gastrointestinal (GI) tract has progressively attracted attention due to the impact of gut-derived hormones on body weight loss and the association of microbiome dysbiosis and host health^1^. The GI tract is a highly dynamic organ system that integrates dietary, microbial, and physical cues to maintain tissue homeostasis and host health^2^. Its primary functions include nutrient digestion and absorption, while simultaneously acting as a selective barrier that limits the passage of harmful antigens and microorganisms^3^. Remarkably, the GI tract adapts rapidly to changes in energy availability and demand, modulating epithelial turnover, barrier function, and mucosal immune responses^4–6^.

Fasting regimes, including calorie restriction and intermittent fasting, have been shown to improve tissue regeneration, health, and lifespan across multiple species^7–11^. Moreover, fasting remodels the intestinal ecosystem. In mice, short and acute fasting enhances the regeneration capacity of intestinal stem cells^12^, which is essential for continuous epithelial renewal. Furthermore, fasting leads to changes in immune function and decreased inflammation^13^. Recent research has shown that many of the fasting-induced intestinal adaptations on tissue regeneration are in fact taking place during the post-fasting refeeding period^14^, highlighting the dynamic nature of nutrient-driven intestinal remodeling. Diet composition also shapes intestinal physiology; for example, it has been shown pre-clinically that a fiber-deprived diet can lead to mucus layer thinning due to a switch in microbial metabolism^15^, while low-energy-density diets in humans are associated with reduced energy intake and improved long-term body weight maintenance in humans^16^. As with dietary interventions, physical activity places metabolic and functional demands on the GI tract, which may elicit acute and chronic adaptations, including improvements in gut motility and alterations in the gut microbiome^17^. Moderate to vigorous exercise can transiently increase epithelial permeability and induce local gut inflammation^5^, although the effects of chronic voluntary exercise on intestinal function remain poorly understood^18^.

Beyond structural and functional adaptations, both diet and exercise modulate the intestinal immune landscape^5,19,20^. The intestine represents the largest immune compartment in the body^21^, and immune cell composition and activity vary along the small intestine. Regional differences in immune specialization are crucial for maintaining mucosal homeostasis^6^, yet they are often overlooked in studies of energy balance and intestinal adaptation^21^. Given the functional differences along the small intestine^21,22^, it is essential to understand how dietary interventions and voluntary exercise influence segment-specific small intestinal immune responses. Here, we used plasma biomarkers and an integrative approach combining intestinal bulk transcriptomic analysis, gene expression analysis, immunohistochemistry, and quantification of secretory epithelial cells (goblet and Paneth cells) to investigate how voluntary exercise alone and in combination with caloric-dilution-diet and fasting-refeeding induce segment-specific adaptations in the proximal, mid, and distal small intestine of mice. To further translate these findings, we analyzed plasma samples from lean and obese individuals before and after an acute exercise bout, enabling a comparative perspective on systemic responses across species.

## 2. Methods and material

### Ethical considerations

Animal studies were conducted with permission from the Danish National Committee for Animal Research (2020-15-0201-00599), from the local animal use committee (SUND, EMED, P22-205, P24-183) and in accordance with the guidelines of Danish legislation governing animal experiments. Human plasma samples were accessed from a previously published study^23^. The study was approved by the ethics committee of Copenhagen and Frederiksberg Communities, Denmark, reported to the Danish Data Protection (P-2019-166), registered at ClinicalTrials.gov (NCT03967691), and performed according to the Declaration of Helsinki. All participants gave written informed consent.

### Animals

Male C57BL/6NRJ mice were purchased from Janvier (Saint Berthevin Cedex, France). The mice were single housed in a temperature (22±2°C) and humidity controlled (55%) room and kept in a 12:12h light/dark cycle with a preceding 30 min half-light period. The mice were fed *ad-libitum* with free access to drinking water. Body composition was assessed using EchoMRI Body Composition Analyzer. Environmental enrichment included nesting, bedding material, small plastic house and cardboard tube.

Specific datapoints from voluntary wheel running protocol were excluded due to measurement errors in both sedentary and running mice. Several datapoints were also missed owing to technical problems with the counters. Investigators (C.B-L. and P.S.) were aware of group allocations in different experiments.

### Experimental setup and diet

#### Study 1: The effects of voluntary wheel running on intestinal gene expression in *ad libitum* chow -fed mice

Male C57BL/6NRJ mice arrived at 7 weeks old. Upon arrival the mice were single housed in a cage with a blocked running wheel. The mice were *ad-libitum* fed a standard chow diet (SAFE D30, Safe Diets, 3.389 kcal/g; 22% protein, 5.3% fat, and 50.8% carbohydrates) with unlimited access to drinking water. After one week, mice were allocated to either an active group (voluntary wheel running) or a sedentary group (blocked running wheel) based on body weight, food intake, and body composition, ensuring comparable baseline measurements between groups. After ∼ 5 weeks of running/sedentary conditions, mice were euthanized just before the start of the dark cycle (ZT12). To minimize the circadian rhythm effect mice from both the sedentary and active group were euthanized in a randomized order and before the lights went off in the animal facility. Tissues (small intestine and colon) were surgically dissected. Tissue used for hematoxylin and eosin staining was collected from a different study in which the same had been handled in the same manner as described above.

#### Study 2: The effects of voluntary wheel running on intestinal gene expression in caloric-diluted fed mice

Male C57BL/6NRJ mice arrived at 4 weeks old. Upon arrival mice were single-housed and fed with control diet (D12450B, 3.8 kcal/g; 20% energy from protein, 10% energy from fat and 70% energy from carbohydrates). After four days and based on the equal distribution of mice regarding body composition and food intake, mice were divided into dietary groups feeding on the control diet or caloric dilution diet in which the food has been diluted with 50% undigestible fiber (D16061505, 1.9 kcal/g; 20% energy from protein, 10% energy from fat and 70% energy from carbohydrates). Food intake, body weight, and body composition were monitored for 15 days. Subsequently, the mice were assigned to either an active (voluntary wheel running) or sedentary group (blocked running wheel) for ∼ 5 weeks. Body weight, food intake, body composition, and running activity were monitored in the following five weeks. On the last day of the experiment, mice were euthanized just before the start of the dark cycle (ZT12). To minimize the circadian rhythm effect mice from both the sedentary and active group were euthanized in a randomized order and before the lights went off in the animal facility. Blood was collected from the trunk into Eppendorf tubes and centrifuged at 3000G for 10 min and then snap-frozen in liquid nitrogen. Tissues (small intestine and colon) were surgically dissected, snap frozen in liquid nitrogen and stored at -80°C before further analysis.

#### Study 3: The effects of voluntary wheel running on intestinal gene expression in fasted and refed mice

Upon arrival, 7-week-old mice were single housed in a cage with a blocked running wheel. The mice were *ad-libitum* fed a standard chow diet (SAFE D30) with unlimited access to drinking water. After one week the mice were divided into two groups: active (EX; voluntary wheel running) or sedentary (SED; blocked running wheel). Body weight, food intake, body composition and running activity were monitored in the following 5 weeks. In the morning (ZT0) on the last day of the experiment before euthanasia the food was removed from all cages and the mice were fasted for 12 hours during daytime. Half of the mice were euthanized in the fasted state at ZT12 (SED-fasted and EX-fasted), the other half was allowed to refeed for one hour before euthanasia (SED-refed and EX-refed). Tissues (small intestine and colon) were surgically dissected and stored based on subsequent analysis.

#### Study 4: The effects of acute treadmill exercise on intestinal length and gene expression analysis in *ad libitum* chow-fed mice

Mice overexpressing the mouse peroxisome proliferative activated receptor, gamma, coactivator 1 alpha under the direction of the mouse muscle creatine kinase promoter (TrC57BL/6-Tg (Ckm-Ppargc1a)31Brsp/J) Jax Strain #:00823. The mice were bred to c57/bl6jr wild-type mice, and the male offspring were singled housed from the age of 10 weeks and after one week used for baseline were split into active (EX; voluntary wheel running) or sedentary (SED; blocked running wheel). The mice were ad-libitum fed a standard chow diet with unlimited access to drinking water. After 43 days, the mice were adapted to the treadmill (TSE systems) for 5 days. On day 1, mice were placed resting on the treadmill for 10 minutes. On day 2: 5 minutes of rest, followed by 5 minutes of walking 5 m/min, slope of 15%. On day 3: 5 minutes of rest, followed by 5 min walking at 5 m/min slope of 15%, followed by 5 min of light running at 10 m/min slope 15%. On day 4: 5 minutes of rest, moderate running for 7 minutes at 15 m/min slope 15%. On day 5: 5 min of rest, graded running speed for 5 minutes starting at 15 m/min ending at 20 m/min, followed by 5 minutes at 20 m/min slope 15%. On day 6, mice were resting in home cage. On the last day of the experiment (day 7) a *rested treadmill group was* resting on the treadmill at the slowest speed 1 m/min for 60 minutes and a *continuous running group was* running for 60 minutes. Continuos running protocol consisted of 5 min at 0 m/min followed by 5 min at m/min, 5 min at 10/min, 5 min at 12m/min, 15 min at 15 m/min, 5 min at 16 m/min, 5 min at 17 m/min, 5 min at 18 m/min and 15 min at 15 m/min all at a 10% inclination. All mice were euthanized immediately after resting/running by quick decapitation. Tissue was surgically dissected, snap frozen in liquid nitrogen and stored at -80°C.

#### Study 5: The effect of acute moderate-intensity exercise training on intestinal inflammation markers in lean and obese individuals

Plasma samples were obtained from a previously published human study^23^. In the original study, lean and obese participants completed an acute exercise bout with or without IL-6 receptor blockade. For the present analysis, only participants who performed the exercise protocol without IL-6 receptor blockade were included (mean men BMI=23.0, *n*=12 and men with obesity BMI=34.3, *n*=9). Stored plasma samples were accessed for secondary analyses under appropriate ethical approval. Two plasma samples per participant were analyzed: one collected immediately before the acute exercise bout, and one collected immediately after exercise completion. Samples had been processed and stored at −80°C according to the protocol described in the original publication and were thawed once for the current analyses.

### Murine tissue harvesting and processing

Mice were euthanized by quick decapitation just before the start of the dark cycle. The small intestine and colon were surgically dissected, and their length was measured and subsequently divided into four segments: proximal (first 2 cm after the stomach), mid (2 cm approximately 15 cm distal of the stomach), distal (last 2 cm before the cecum), and colon (most distal 2 cm before the rectum). All tissues were stored based on future analysis.

### Bulk transcriptomic profiling by RNA sequencing

RNA from proximal, mid, and distal intestinal sections from active and sedentary mice (*study 1*) euthanized during *ad libitum* feeding was obtained using TRIzol method. Briefly, the tissue was homogenized using 1 mL of TRIzol in a Tissuelyser (Qiagen Tissuelyser Retsch MM300). Cell debris was removed by centrifuging the sample at 12,000 g for 10 min at 4 °C. The TRIzol phase was transferred into new 2 mL tubes with 200µL chloroform. After centrifugation, RNA was precipitated using ice-cold isopropyl alcohol and subsequently washed with 75% ethanol and dissolved in Ultrapure RNase/DNase free water. RNA purity was measured using Nanodrop spectrophotometer (nanodrop one, Thermo Fischer Scientific, USA). RNA samples were submitted to BGI for sequencing. Two mid-intestinal sections from sedentary mice were discarded from sequencing given the low-quality of the RNA. The RNA-seq libraries for mid and distal sections were prepared for all high-quality RNA samples using mRNA with poly A tail enrichment whereas stranded RNA-seq libraries for proximal section were sequenced using rRNA-depletion library given low-quality control of RNA. The libraries were sequenced in paired-end 2x100 nt mode on a DNBSEQ platform. We generated ∼5 Gb of RNA-seq data for mid and distal sections and ∼7 Gb of RNA-seq data for each for proximal section. Quality control of the raw sequencing data was performed with SOAPnuke v2.1.5^24^ with default options and the reads for each of the samples were mapped to the *Mus musculus* genome assembly GRCm39 using STAR v2.7.11b^25^. The gene counts were retrieved using featureCounts^26^ v2.0.6 and then built into a matrix. Low-count genes were filtered based on the count-per-million and sample library size (the gene is expressed in at least one replicate). The expression abundance was quantified using cufflinks v2.2.1^27^, retrieving the fragments per kilobase of transcript per million mapped reads (FPKM) values. The expression clustering of the replicate samples for each of the sequenced sections were visualized using PCAs from R package ggplot2^26^. Differential expression analyses were conducted separately for each of the sequenced tissue sections using DESeq2^28^. Volcano plot was used for visualizing each pair-wise comparison of each section using R package ggplot2^29^, dplyr function from R package tidyverse^30^ and ggrepel. Genes with p-value<=0.01 and absolute log_2_FC greater than 1, p-values<=0.05 and absolute log2FC greater than 1 and p-value<=0.01 were highlighted in red, pink, and brown, respectively. Non-significant genes were labelled in grey. We then counted the number of genes with p-value<=0.01 from each pair-wise comparison of each section and highlighted the overlap of genes between the comparisons using R package VennDiagram^31^. The Gene Ontology (GO) functional enrichments of the genes with pvalue <= 0.01 were conducted using enrichGO function of clusterProfiler R package^32^ and the top 10 most significant terms of GO biological process were then visualized. Functional protein association networks were conducted using the Web-based tool STRING^33^ and the proteins belonging to the biological processes with the five highest false discovery rates were highlighted.

### Gene expression analysis

Gene expression profiling in the small intestine (proximal, mid and distal segment) was conducted both in sedentary and active mice. Total RNA was isolated as described above. RNA was converted into cDNA using cDNA Reverse Transcription Kit (Cat# 4368813, Life Technologies) following the manufacturer’s instructions. Quantitative PCR (qPCR) were performed with PowerUpTM SYBRTM Green Master Mix (Cat# A25780, ThermoFisher Scientific) and 300 nM of forward and reverse primers and using ViiA 7 Real Time PCR system (Applied Biosystems) for amplification. *Gadph* was used as housekeeping gene and used to quantify the relative gene expression amount. Primers were synthetized by TAG Copenhagen.

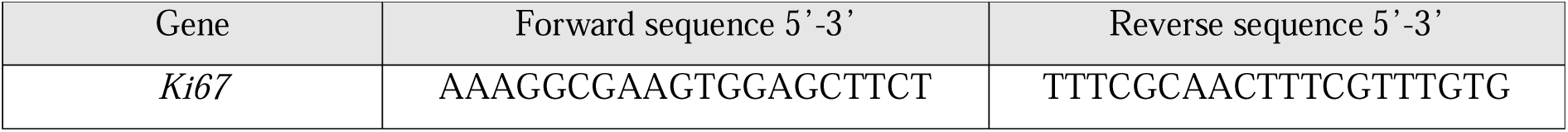
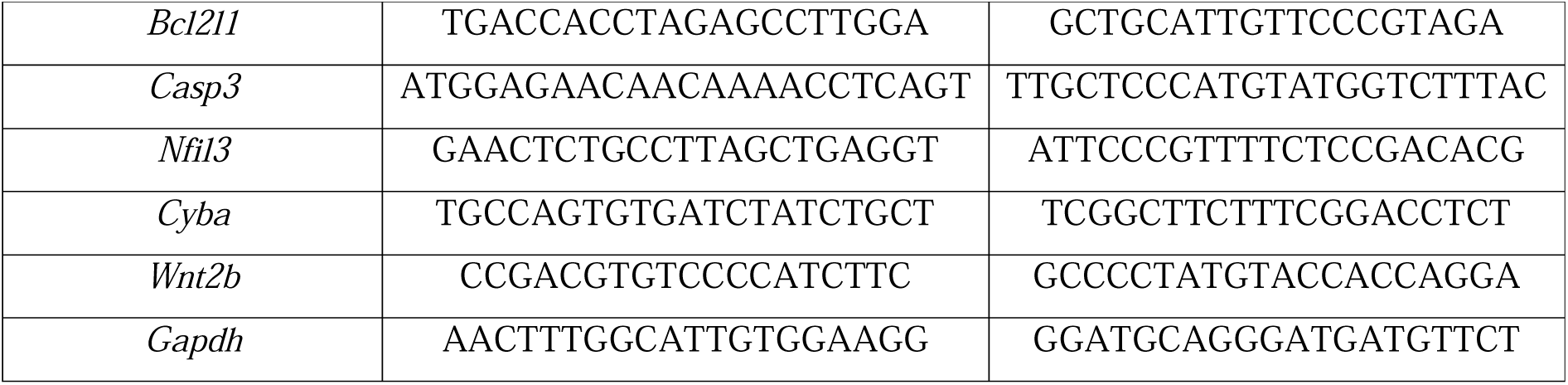

### Immunostaining and image analysis of small intestine

The small intestine was surgically removed and divided into segments (proximal, mid and distal) and placed on 4% PFA until mounting each segment in paraffin blocks. The samples were cut in slide sections of 4 µm and dewaxed through Tissue clear, alcohol and tap water. To retrieve antigens, Ki67 and Cd8a sections were placed in a TEG-buffer pH=9 and Cd4 were placed in Citrat buffer pH=6 and boiled in a microwave oven for 15 minutes. Next, 10 minutes pre-incubation in 2% BSA was performed ahead of overnight incubation at 4°C with a primary antibody. Specific dilutions were used for each antibody: 1:500 for Cd4 (Abcam,83685), 1:250 for Cd8a (cellSignaling technology, D4W2Z) and 1:400 for Ki67 (Abcam, ab16667). The sections were incubated for 40 minutes with a second layer of antibodies to amplify the reaction. Diluted 1:200 biotinylated secondary immunoglobulins were used (goat anti-rabbit: Vector Laboratories, BA-1000 and BA-2000). Next, hydrogenperoxide 3% was added to block endogenous peroxidase. The third layer consisted of a preformed avidin and biotinylated horseradish peroxidase macromolecular complex (Elite ABC; Vector Laboratories, PK-6100) and incubated for 30 minutes. The reaction was developed by the use of ImmPACT DAB code nr:SK-4105 for 15 minutes, Finally, counterstaining with Mayers Hemalun (Merck Eurolab, Darmstadt, Germany) was performed.

### Hematoxylin and eosin staining

Intestinal tissue was surgically dissected, washed in PBS and placed in 4% PFA for 24-h at RT. Samples were stored in 4% PFA at 4L and then embedded in paraffin and cut by In-Lab (Denmark, https://in-lab.dk/). HE-staining was used to quantify goblet cells and Paneth cells. Goblet cells were identified by their mucin-filled cytoplasm and basally displaced nuclei within the intestinal epithelium, whereas Paneth cells were identified by their characteristic eosinophilic granules located at the base of intestinal crypts. Histological images were acquired using EVOS5000 light microscopy. Villus height and crypt depth was measured manually with Fiji ImageJ. Quantification was performed in well-oriented crypt–villus units. Goblet cells were counted along the villus epithelium and expressed as the number of cells per villus, while Paneth cells were quantified as the number of cells per crypt. Three sections per intestinal segment from each mouse were analyzed. All villus and crypts were measured. Quantification was performed in a blind manner.

### Mouse serum analysis

Mouse serum concentrations of soluble CD14 and intestinal fatty acid–binding protein (I-FABP/FABP2) were quantified using commercially available enzyme-linked immunosorbent assay (ELISA) kits according to the manufacturers’ instructions. CD14 levels were measured using the Mouse CD14 ELISA Kit (Abcam, ab242238), and I-FABP levels were measured using the Mouse FABP2/I-FABP Colorimetric ELISA Kit (Novus Biologicals, NBP2-82214). Serum samples were assayed at two different dilutions (I-FABP: 1:40 and 1:80 and CD14: 1:5) and measured in singlet. Absorbance was measured using a microplate reader at the specified wavelengths, and concentrations were calculated from standard curves generated using supplied recombinant standards.

### Human plasma analysis

Plasma concentrations of intestinal fatty acid–binding protein (I-FABP/FABP2), CD14, interleukin-32 (IL-32), and regenerating islet–derived protein-3 alpha (Reg-3α) were quantified using electrochemiluminescence-based U-PLEX immunoassays (Meso Scale Discovery, MSD) according to the manufacturer’s instructions. Plasma samples were assayed at the recommended dilutions. Signals were detected using an MSD MESO Quick Plex SQ 120 and the data were analyzed using The DISCOVERY WORKBENCH 4.0 Analysis Software and concentrations were calculated from standard curves generated with the supplied calibrators. Only values within the linear range of the assays were included in the analysis.

### Data presentation and statistical analysis

The graphs were made using GraphPad prism 9 (GraphPad, La Jolla, USA). All data are presented as mean±SEM. The number of animals included in each analysis is indicated in the corresponding figure legends, and missing values reflect samples that were excluded due to technical errors during sample processing or assay execution. For murine metabolic and intestinal length analysis, GraphPad prism 9 (GraphPad, La Jolla, USA) was used using two-way ANOVA. For gene expression analyses, two-way ANOVA was conducted performed using IBM SPSS Statistics (version 29.0.1.0, IBM Corp., Armonk, NY, USA). If Shapiro-Wilk Normality test and/or Levene’s test for homogeneity of variances were significant, a Generalized Linear Model (GzLM) was used. Statistical analysis employed is noted in the figure legends. P-values<0.05 was considered significant. *P<0.05, **P<0.01, ***P<0.001.

## 3. Results

### 3.1 Active mice fed ad libitum chow exhibit intestinal segment-specific transcriptomic and secretory epithelial cell adaptations

To investigate how voluntary wheel running influences intestinal epithelial and molecular function, we conducted bulk-RNAseq in the proximal, mid and distal small intestine from sedentary and active mice (Figure 1A). Most changes were observed in the proximal small intestine, where 464 genes were affected by physical activity. In the mid and distal small intestine, the number of genes were 54 and 45, respectively (Figure 1B and Supp. Figure 1A-C). Some genes affected by physical activity overlapped between intestinal segments. The expression of *Ndrg1*, *Gvin2*, *Heatr9*, *Tcaf2*, *Ptgr1*, *Ces1e*, *Pim3*, and *Clic4* were affected by voluntary wheel running in both the proximal and mid regions. Notably, these genes are all associated with immune responses or epithelial stress pathways. *Nfil3* and *Cyba* were affected by voluntary wheel running in both proximal and distal regions, and *Mt2* was affected by voluntary wheel running in both the mid and distal regions (Figure 1B). Next, we assessed the function of differential expressed genes in each intestinal segment. In the proximal small intestine, the expression of the genes affected by physical activity were mostly downregulated, and most genes were related to immune system functions (Figure 1C, Supp. Figure 2A). In the middle segment, the genes that were significantly regulated by physical activity were associated with muscle organ development as well as processes involved in cellular ketone and steroid metabolism (Figure 1D and Supp. Figure 2B). The expression of genes: *Rdh7, Rdh16f2,* and *Retsat*, which are genes encoding enzymes related to vitamin A processing critical for immune homeostasis ^34,35^, were upregulated by physical activity (Figure 1D, Supp. Figure 2B). In the distal small intestine, 45 genes were affected by physical activity, most of which were immune-related, mirroring the gene pattern detected in the proximal small intestine (Figure 1E, Supp. Figure 2C). Interestingly, *Cyba, Il1b, Lbp* were downregulated by physical activity, and these genes are related in reducing signaling inputs that normally activate NF-κB^36,37^, resulting in a lower inflammatory transcriptional state and a lower basal drive for IL-6 production. *Cyba* was also downregulated by physical activity in the proximal segment. Whereas *Nfil3*, a transcription factor required for the development of gut-associated ILC3s^38^, was downregulated in both proximal and distal segments following physical activity, suggesting a reduction in basal type 3 immune programming. In contrast, *Mt2*, a cytoprotective and antioxidant gene involved in immune modulation, was upregulated, consistent with enhanced epithelial stress resilience^39^. Together, these changes indicate a segment-specific shift toward reduced inflammatory tone and improved mucosal homeostasis in response to physical activity.

**Figure 1.**
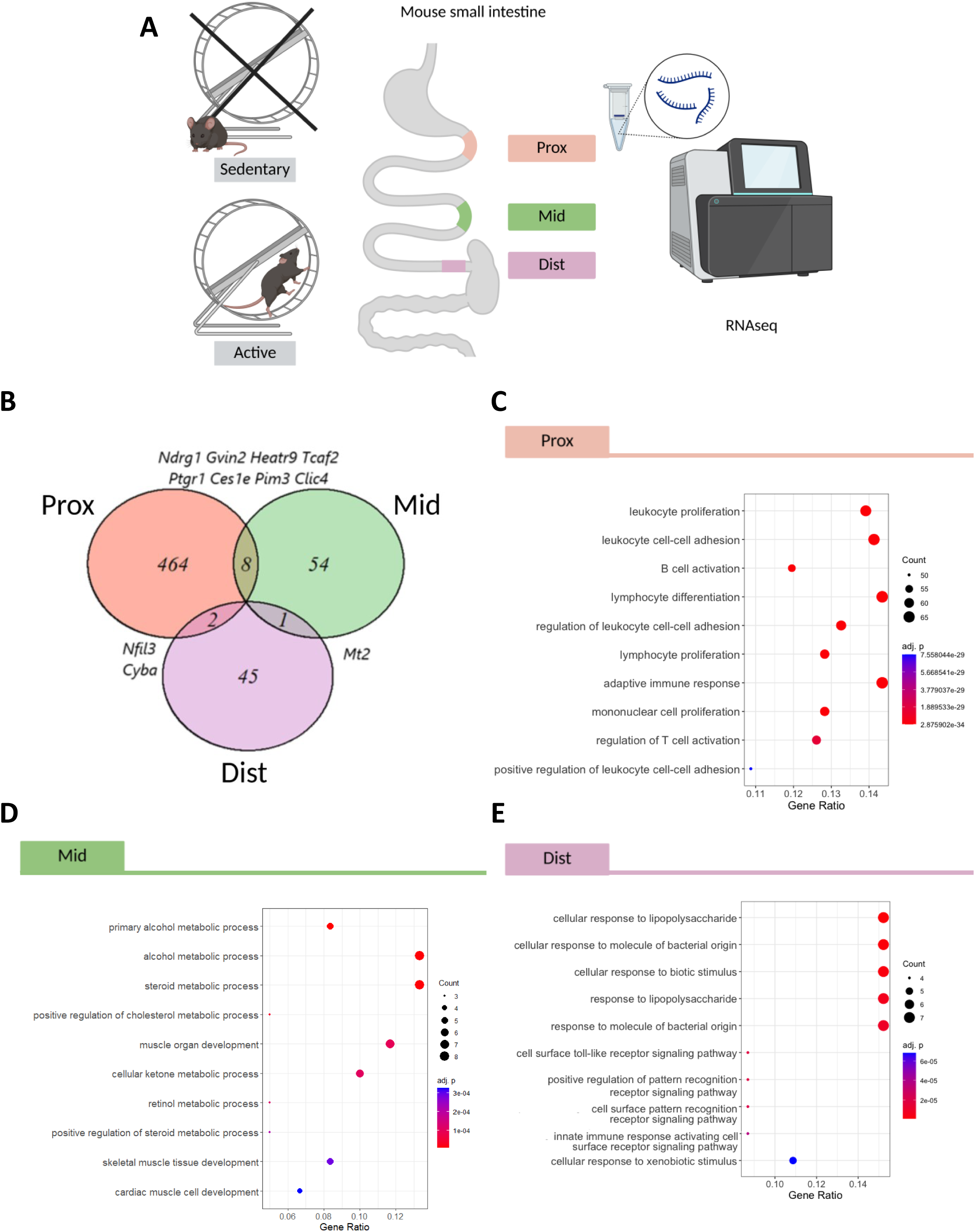
Voluntary wheel running under *ad libitum* chow feeding conditions induces segment-specific transcriptomic and secretory epithelial cell adaptations. A) Schematic overview of study setup with color scheme. B) Venn diagram of three intestinal segments. C) Top 10 most significant terms of GO biological processes for the proximal segment. D) Top 10 most significant terms of GO biological processes for the middle segment. E) Top 10 most significant terms of GO biological processes for the distal segment. A was created using BioRender.

**Figure 2.**
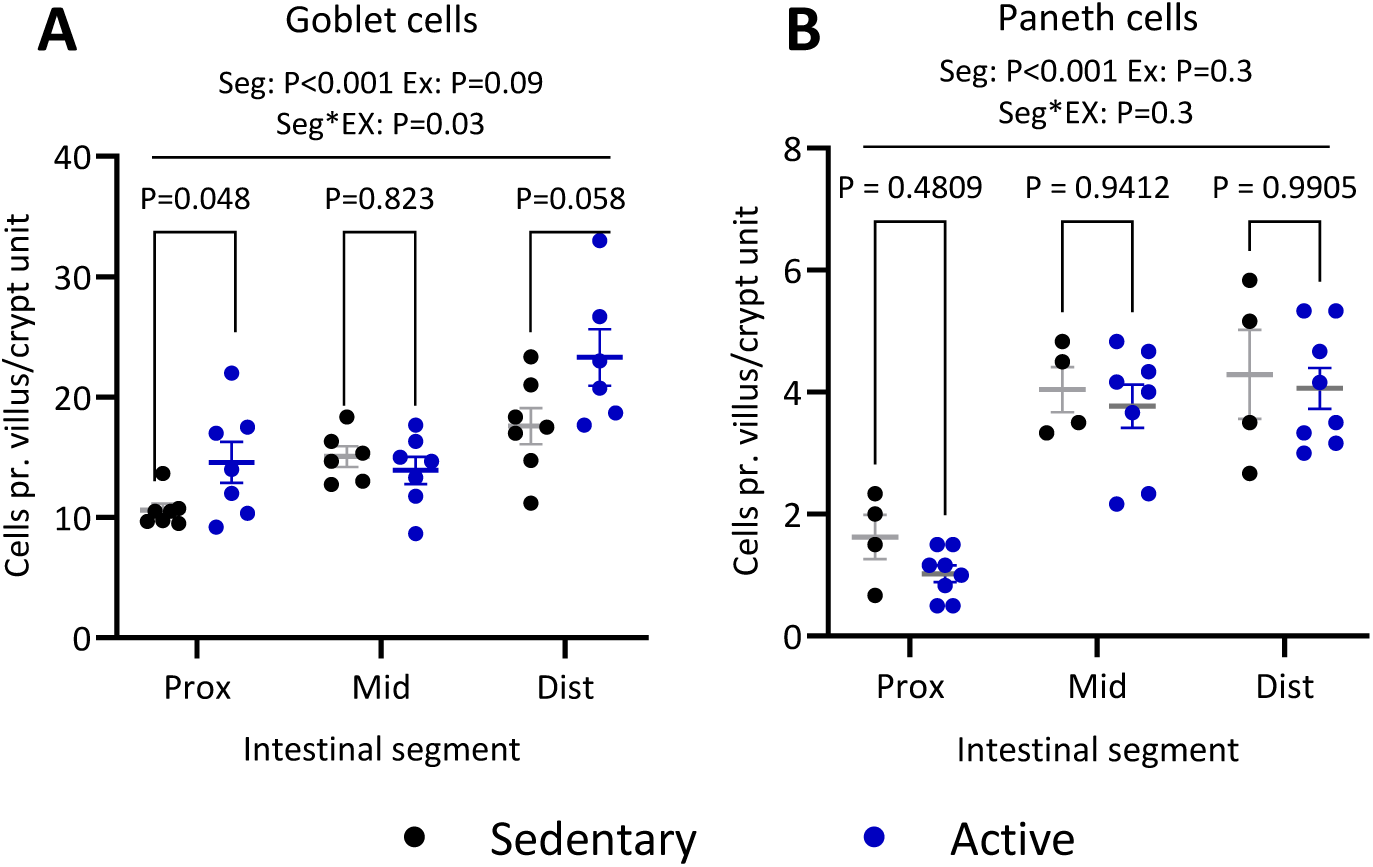
Voluntary wheel running under ad libitum chow feeding conditions has a modest impact on the number of small intestinal Goblet and Paneth secretory epithelial cells. A) Number of goblet cell/villus in the three segments of the small intestine in sedentary (black) and active (blue) mice. B) Number of Paneth cell/villus in the three segments of the small intestine in sedentary (black) and active (blue) mice. Two-way ANOVA was used to compare the effects of exercise and segment, and post hoc multiple comparisons were performed using Bonferroni’s correction. Data is shown as mean ± SEM.

To complement this data, we quantified secretory epithelial cells, namely goblet and Paneth cells in sedentary and active mice. Both cell types contribute to mucosal barrier function, producing mucus and antimicrobial peptides, respectively, and their regulation may reflect adaptive epithelial responses to metabolic and physical challenges. The number of goblet cells increased along the length of the small intestine, aligning with the literature^40^, and trended towards being induced by physical activity (mean difference: 2.8±1.5 cells/villus CI: [0.5 to 6.1], P=0.09) (Figure 2A). However, goblet cell number was significantly greater in the proximal small intestine of active mice compared to sedentary (mean difference: 4.0±1.8 cells/villus, P=0.048) and tended to be enhanced in the distal small intestine (mean difference: 5.7±2.7 cells/villus CI: [-0.24 to 11.65], P=0.058). We did not observe a difference in the number of Paneth cells between the two groups in any of the analyzed intestinal segments, however Paneth cell number increased from the proximal to the distal small intestine (Figure 2B). Taken together, our data suggests that regular voluntary wheel running not only reprograms molecular pathways but might also remodel key barrier-supporting cell populations in a segment-specific manner.

### 3.2 Chronic caloric dilution feeding modulates intestinal molecular signature and plasma markers in both sedentary and active mice

Calorie restriction is thought to increase health, lifespan, and tissue-regeneration^7,10,11^. Introducing a caloric dilution diet to mice enables calorie restriction without inducing the hunger symptoms observed during regular calorie restriction^41,42^. To determine the effect of combined exercise and caloric dilution intervention we monitored mice for six weeks. 4-week-old male C57BL/6J mice were fed either a caloric diluted (CD) or a control (CT) diet and then allocated a working (EX) or a blocked (SED) running wheel for five weeks (Figure 3A). The caloric diluted diet is 50% kcal/g compared to the control diet. Previously published results on running distance, energy balance and body composition from this cohort^43^ were integrated with newly collected small intestinal data to assess how the small intestine responds to the combination of caloric dilution and exercise. In summary, after 5 weeks of running, there was no difference in running distance between EX-CT or EX-CD (Supp. Figure 3A) and CD fed mice consumed 80.5±3.6 g more food compared to CT fed mice. Despite increased dietary intake, caloric dilution resulted in decreased energy intake of 22.3±9.1 kcal over the period (Supp. Figure 3C). Similarly, active mice consumed 15.9±1.4 g more food than their sedentary counterparts (Supp. Figure 3B). There was no significant effect of either caloric dilution or exercise on five-week body weight change (Supp. Figure 3D). However, exercise, but not CD, positively affected body composition by increasing lean mass by 3.6±1.7% (Supp. Figure 3E) and decreasing fat mass by 43.8±13% (Supp. Figure 3F). Increased physical activity augments food intake which drives a lengthening of the small intestine and colon^44^. In this cohort, the CD diet fed mice consumed more food daily than the CT diet fed mice possibly resulting in increased intestinal growth. Caloric dilution fed mice had on average 5±0.8 cm longer small intestine (Figure 3B), and 2.8±0.2 cm longer colon (Figure 3C) than the CT diet fed mice. Running led to a 1.8±0.8 cm increase in small intestine length, and 0.8±0.2 cm increase in colon length compared to sedentary control mice. Small intestine length was positively correlated with total food intake in the active mice independent of diet (EX-CT mean slope: 0.038 CI: [0.02 to 0.06] P=0.004, EX-CD mean slope: 0.05 CI: [-0.001 to 0.09] P=0.055), but not in the sedentary controls (SED-CT P=0.81, SED-CD P=0.79) (Figure 3D), substantiating our previously reported findings^44^. Intestinal length tended to be correlated to total running distance (Figure 3E) (EX-CT mean slope: 0.007 CI: [-0.003 to 0.02] P=0.13, EX-CD mean slope: 0.014 CI: [-0.003 to 0.03] P=0.086). These data reveal that exercise-induced small intestinal growth is associated with the exercise-induced increase in food intake.

**Figure 3.**
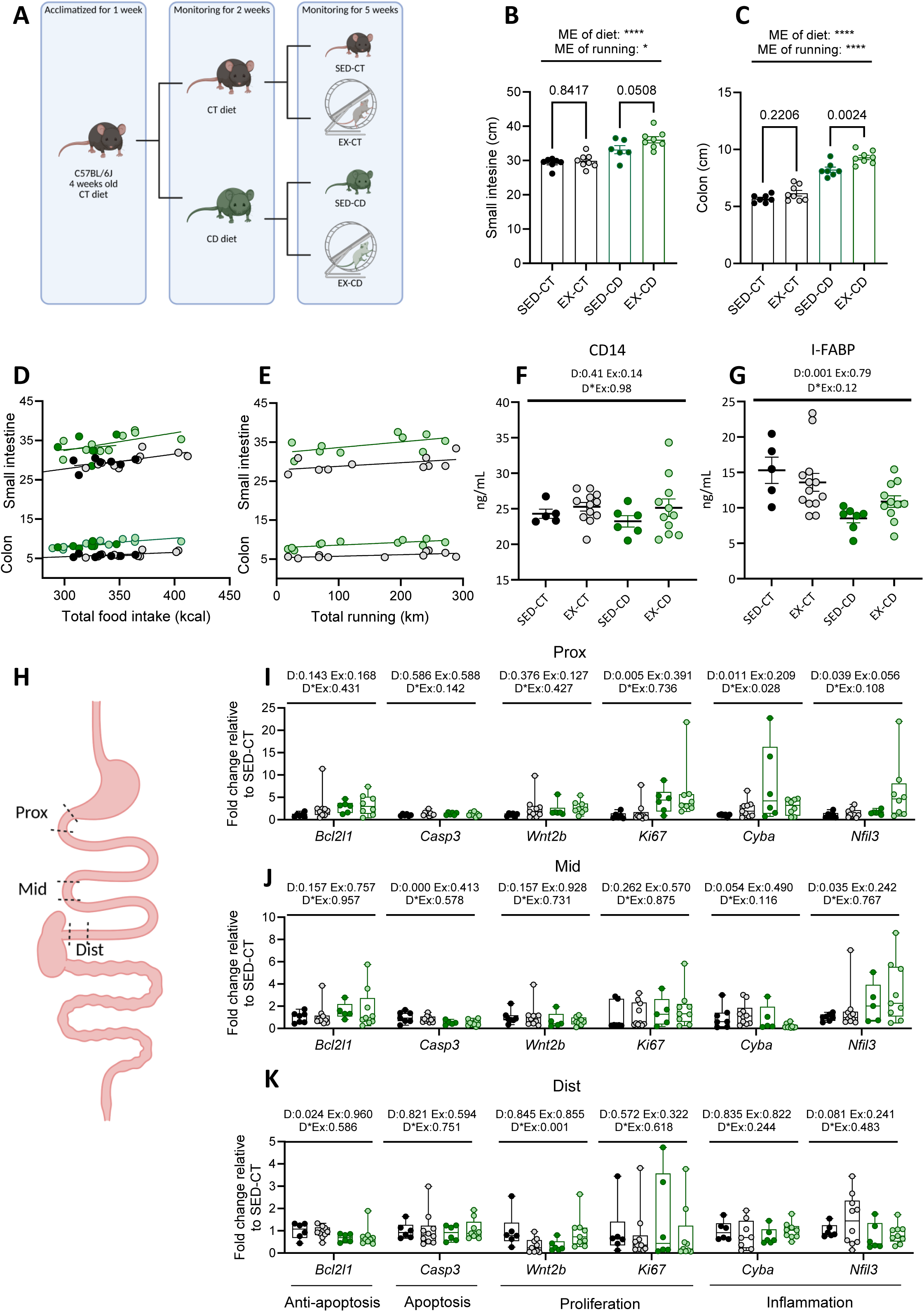
Chronic caloric dilution feeding modulates intestinal molecular signature and plasma markers in both sedentary and active mice. A) Schematic overview of study setup with color scheme. B) Small intestine length in *ad-libitum* fed mice. C) Colon length in *ad-libitum* fed mice. Intestinal length is compared by two-way ANOVA with Bonferroni multiple comparison. D) Simple linear regression analysis of total food intake (kcal) in relation to small intestine and colon length. E) Simple linear regression analysis of total running distance (km) in relation to small intestine and colon length. F) Plasma concentration of CD14 and G) I-FABP compared by two-way ANOVA. H) Illustration of the intestinal segments used for analysis (prox, mid and dist). I) Intestinal gene expression in the proximal small intestine, J) mid small intestine and K) distal small intestine in the four groups. Two-way ANOVA was used for evaluating the effects of calorie-diet intervention (D), voluntary wheel running (Ex) and its interaction (D*Ex), if normality and/or homogeneity of variance were violated, GzLM was used. Data is shown as mean± SEM and box plots show the median, interquartile range, and range of the data with individual mice represented as dots. A and H were created using BioRender.

Measurement of plasma markers further supported differential systemic responses. Circulating CD14, an indicator of monocyte activation and potential gut-derived inflammatory signaling^45^, remained unchanged by exercise or dietary intervention (mean difference: 1.9±1.2 ng/mL CI: [-0.7 to 4.5], P=0.14) (Figure 3F).

However, I-FABP, an epithelial-derived protein released into the circulation upon small intestinal epithelial damage^46^, was lower in mice fed the CD diet and was not affected by voluntary wheel running (Figure 3G). To further investigate the independent and combined effects of caloric dilution and voluntary wheel running, we performed targeted qPCR analysis of genes related to epithelial turnover and immune regulation in distinct segments of the small intestine (Figure 3H). The analysis revealed segment-specific effects of diet, whereas no clear effects of physical activity were detected in this targeted panel. In the proximal small intestine, dilution feeding increased expression of *Ki67* (proliferation marker)*, Cyba* (marker immune activity), and *Nfil3* (promotes cell survival) (Figure 3I), suggesting enhanced epithelial turnover and modulation of immune-related signaling under reduced caloric density. In the mid small intestine, caloric dilution reduced expression of *Caspase-3* (a marker of apoptosis) and increased expression of *Nfil3* (Figure 3J), consistent with region-specific alterations in apoptotic and immune-associated pathways. In the distal small intestine (Figure 3K), *Bcl2l1* (a marker of anti-apoptosis) expression was reduced by the dilution diet, whereas in the proximal segment a non-significant trend toward increased *Bcl2l1* expression was observed, further supporting spatially distinct regulation of epithelial survival pathways.

In summary, voluntary wheel running modestly affected the expression of the analyzed genes across all three intestinal regions under control and caloric-dilution diet. This contrasts with the bulk transcriptomic analysis performed here in the three intestinal segments of sedentary and active mice fed a chow diet, suggesting that exercise-induced transcriptional adaptations may depend on diet composition.

### 3.3 Sedentary and active mice displayed segment-specific intestinal transcriptional responses to fasting and refeeding

To investigate how acute nutritional challenges interact with chronic exercise, we subjected sedentary and active mice to 12 hours of daytime fasting (06:00–18:00). Half of the mice in each group were subsequently allowed to refeed *ad libitum* for 1 hour (Figure 4A). Active mice had longer small intestine than sedentary mice^44^, and there was a trend towards a positive correlation between small intestine length and total calorie intake in active but not sedentary fasted mice (Figure 4B). Given the role of physical activity on intestinal immune genes (Figure 1), we next quantified the number of CD8 (Figure 4C), Ki67 (Figure 4D) and CD4 (Figure 4E) positive cells in the three small intestinal segments (Figure 4F) to investigate mucosal immune cell composition and proliferative status. The amount of CD4, CD8 and Ki67 cells differentiated between segments. Accordingly, CD4 and CD8 were most abundant in the proximal small intestine, whereas Ki67 positive cell numbers were highest in the distal segments (P = <0.001, <0.001, and <0.01, respectively). There was no significant effect of physical activity on the number of these cells in any region, although there were modest tendencies towards a reduced number of CD4 and Ki67 positive cells with physical activity (mean difference: 1.8±1.1 cells/villus CI: [-0.4 to 4.0], P=0.11 and 2.3±1.3 cells/villus CI: [-0.4 to 4.9], P=0.088, respectively).

**Figure 4.**
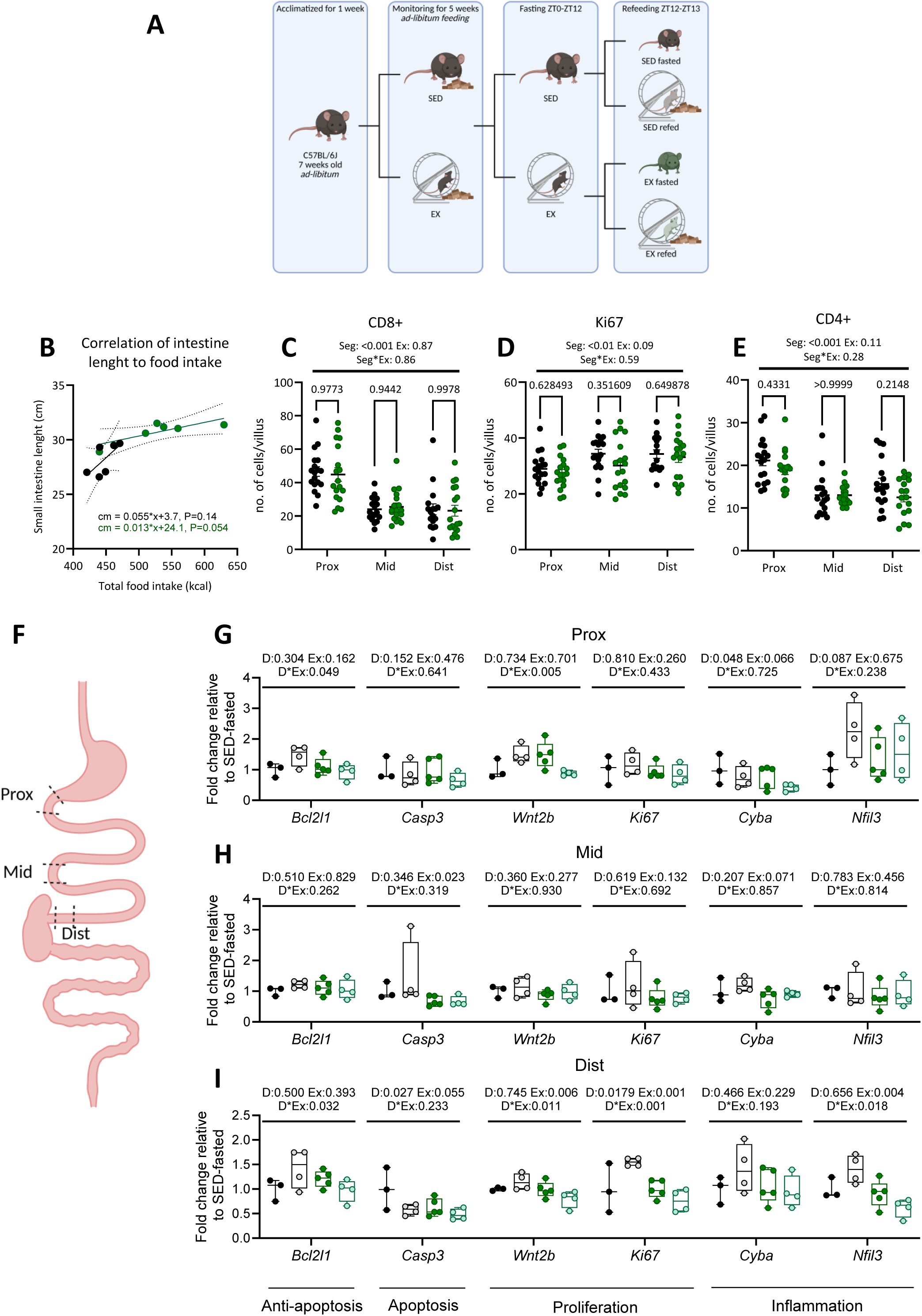
**Sedentary and active mice display to segment-specific intestinal transcriptional responses to fasting and refeeding**. A) Schematic overview of study setup with color scheme. B) Simple linear regression analysis of total food intake (kcal) in relation to small intestine length in fasted sedentary (bold black) and active (bold green) mice. Small intestinal C) CD8L cells, D) Ki67L cells and E) CD4L cells compared by two-way ANOVA with Bonferroni multiple comparison. F) Illustration of the intestinal segments analyzed (prox, mid and dist). Intestinal gene expression in the G) proximal, H) mid and I) distal small intestine. Two-way ANOVA was used for evaluating the effects of refeeding (D), voluntary wheel running (Ex) and its interaction (D*Ex), if normality and/or homogeneity of variance were violated, GzLM was used. Data is shown as mean± SEM and box plots show the median, interquartile range, and range of the data with individual mice represented as dots. A and F were created using BioRender.

In the same groups of mice, we performed targeted qPCR analysis of genes involved in apoptosis, proliferation, epithelial turnover, and immune regulation which revealed segment-specific effects in response to refeeding. In sedentary mice, refeeding induced significant upregulation of *Bcl2l1, Wnt2b,* and *Nfil3* in the proximal small intestine (Figure 4G), while the distal small intestine showed increased expression of *Bcl2l1, Wnt2b, Cyba*, and *Nfil3* (Figure 4I). The mid small intestine exhibited minimal transcriptional response (Figure 4H). In contrast to the effect of refeeding in sedentary mice, chronically active mice did not exhibit the same refeeding-induced upregulation. Instead, a trend toward downregulation of *Wnt2b* was observed in proximal and distal segments, suggesting that chronic exercise dampens acute transcriptional activation of apoptotic, proliferative, and immune-related pathways. This dampening may represent a protective adaptation, limiting excessive epithelial turnover and immune signaling in response to nutrient intake, thereby helping to maintain intestinal homeostasis, preserve barrier function, and reduce low-grade inflammation. Such adaptations likely contribute to the segment-specific remodeling of the intestine observed with chronic exercise, supporting more stable epithelial and immune function in physically active mice.

### 3.4 Acute exercise elicits transient intestinal and immune responses in both mice and humans

To assess the systemic impact of acute exercise on intestinal and immune-related plasma markers, we first performed an acute exercise intervention in sedentary and active mice. By combining acute treadmill exercise with a 5-week voluntary exercise regimen (Figure 5A), we assessed segment-specific intestinal morphology and targeted gene expression to determine how chronic and acute physical activity interact at the tissue and molecular levels. Since the genotype of the mice did not affect the analysis, only types of exercise were considered for the analysis. Small intestinal and colonic morphology showed distinct responses to chronic and acute activity. Consistent with previous findings, chronically active mice had longer small and large intestines than sedentary control mice, likely reflecting increased food intake. Active mice had 2.0±0.9 cm (P=0.046) longer small intestine and 0.47±0.17 cm (P=0.01) longer colon than sedentary controls (Figure 5B-C). Interestingly, acute exercise caused a transient shortening of the small intestine (mean difference: - 2.1±0.9 cm, P=0.038) but not colon (mean difference -0.08 cm CI: [-0.4 to 0.2], P=0.62), indicating that acute exercise predominantly influences small intestinal morphology.

**Figure 5.**
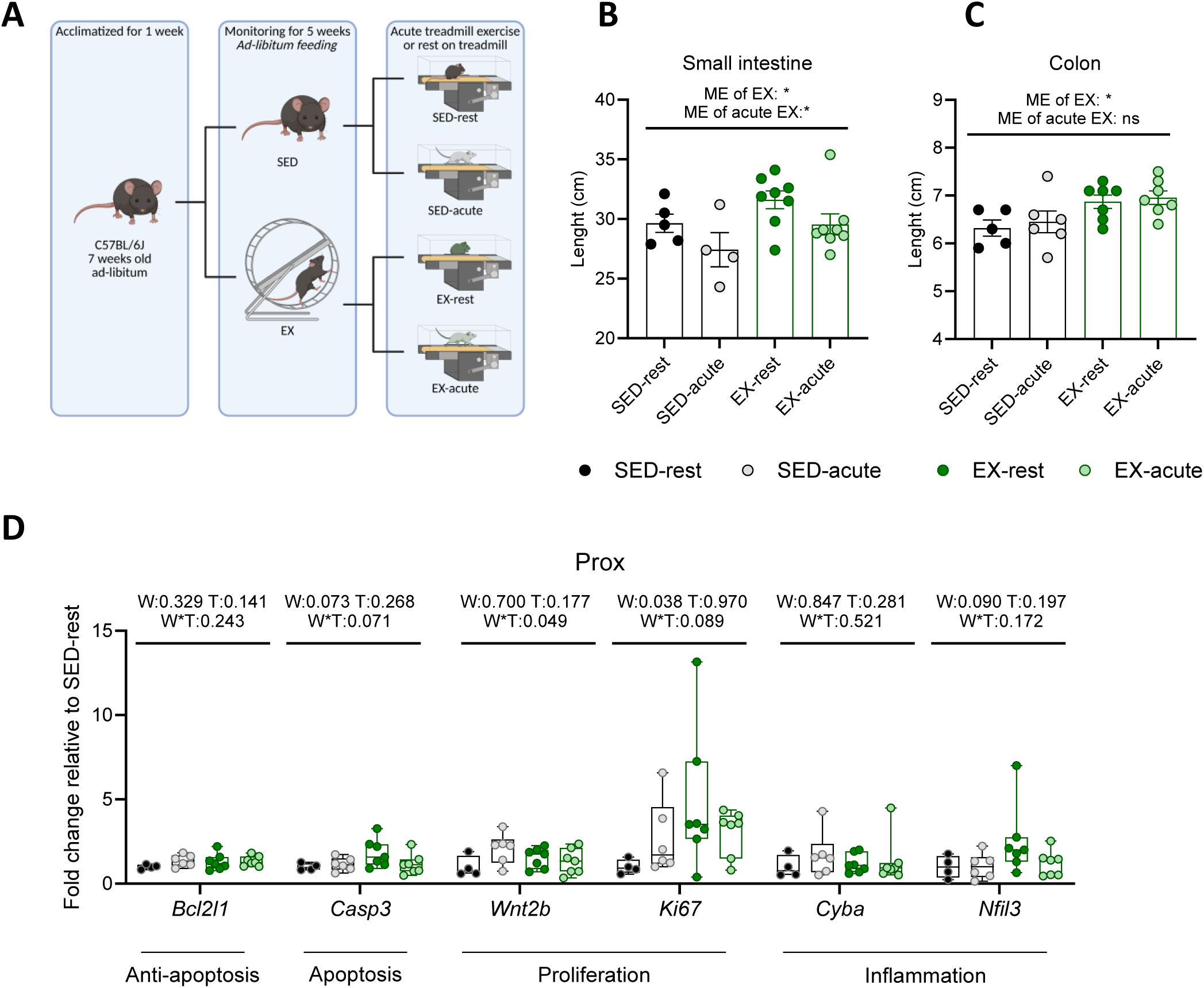
**Acute exercise induces a transient shortening of the small intestine and activates intestinal stress and regenerative pathways in sedentary mice, which is blunted by chronic voluntary activity**. A) Schematic overview of study setup with color scheme. B) Small intestine and C) Colon length compared by two-way ANOVA. D) Intestinal gene expression in the proximal small intestine. Two-way ANOVA was used for evaluating the effects of voluntary wheel running (W), acute training (T), and its interaction (T*W), if normality and/or homogeneity of variance were violated, GzLM was used. Data is shown as mean± SEM and box plots show the median, interquartile range, and range of the data with individual mice represented as dots. A was created using BioRender.

At the molecular level, targeted qPCR analysis of genes involved in apoptosis, proliferation, epithelial turnover, and immune regulation revealed complementary patterns in the proximal segment. In particular, acute exercise induced upregulation of *Bcl2l1, Casp3, Wnt2b,* and *Ki67* in sedentary mice, reflecting a rapid activation of apoptotic and proliferative pathways in response to acute stress. In chronically active mice, these acute responses were blunted, suggesting that repeated exercise leads to adaptive remodeling of the small intestine. Chronic physical activity itself was associated with elevated baseline expressions of *Casp3, Ki67,* and *Nfil3,* indicating persistent modulation of apoptotic, proliferative, and immune-related signaling (Figure 5D). Together, these findings show that acute exercise transiently activates intestinal stress and regenerative pathways in sedentary mice, whereas chronic voluntary activity blunts these acute responses while maintaining elevated basal gene expression, consistent with adaptive remodeling of the intestinal epithelium Next, we measured plasma concentrations of CD14, IL-32, Reg-3α, and I-FABP, as markers of immune activation and intestinal permeability, in lean and obese human participants before and immediately after a single bout of moderate-intensity exercise (Figure 6A). In both lean individuals and in individuals with obesity ^23^, plasma CD14 increased significantly following acute exercise (65953±23361 pg/mL) (Figure 6B). While baseline CD14 was slightly higher in participants with obesity, body weight alone did not correlate with CD14 levels. We did, however, observe a significant interaction between exercise and weight status, suggesting that the magnitude of exercise-induced CD14 increase differs between lean and obese individuals. IL-32 concentrations were also elevated after acute exercise (572.8±144.8 pg/mL), with no significant effect of obesity or interaction, indicating a robust exercise-induced response across both lean and obese participants (Figure 6C), similarly did Reg-3α show a small but significant increase after acute exercise (0.92±0.34 pg/mL), with no effect of body weight or interaction (Figure 6D). Finally, I-FABP, a marker of small intestinal epithelial stress, increased following acute exercise (589.6±184.6 pg/mL), with no significant influence of body weight or interaction (Figure 6E). These results demonstrate that acute moderate-intensity exercise triggers rapid systemic responses of both immune and intestinal epithelial markers, largely independent of lean and obese conditions.

**Figure 6.**
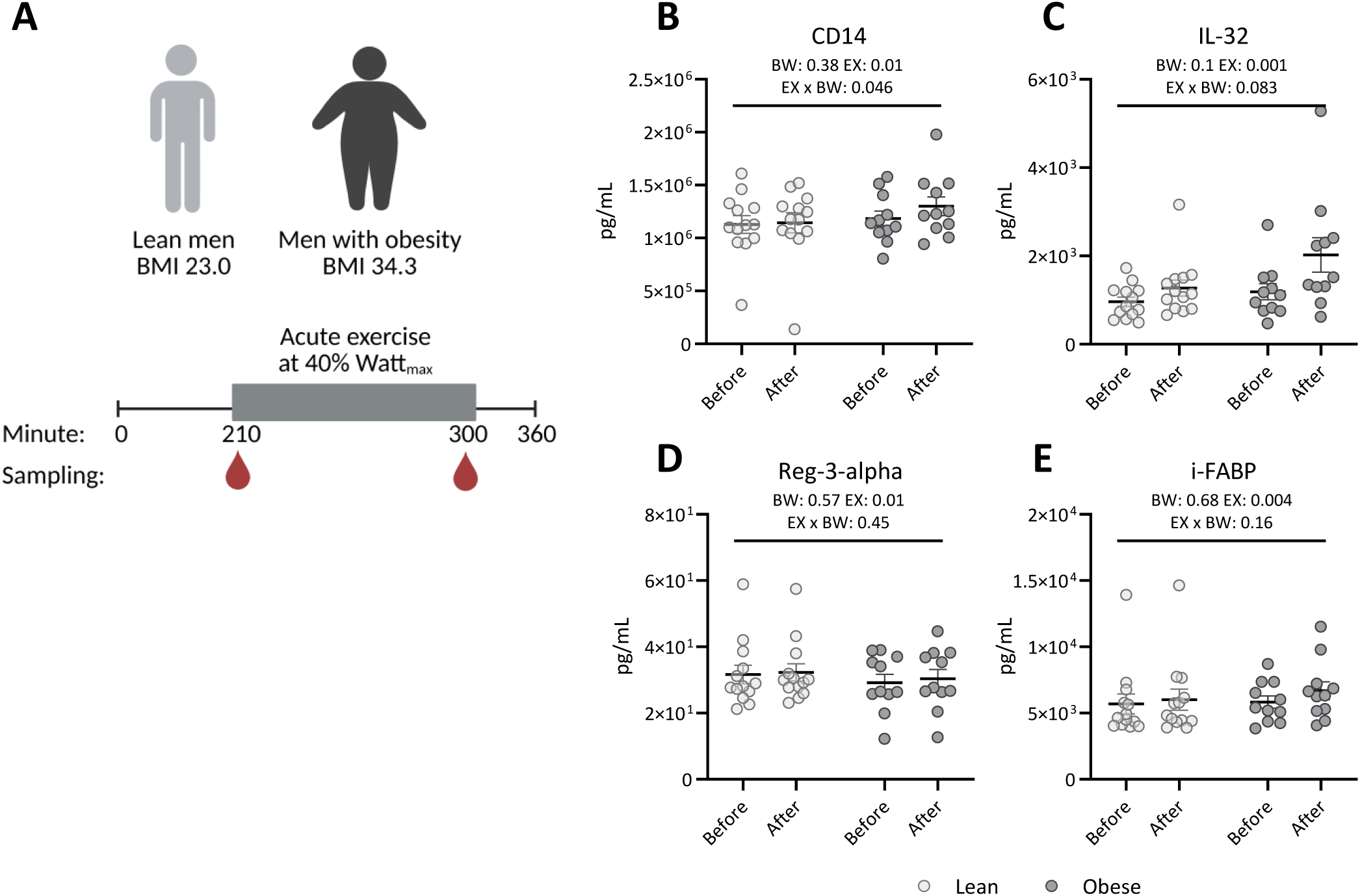
**Acute moderate-intensity exercise induces a systemic response in immune and intestinal epithelial markers in both lean and obese participants**. A) Schematic overview of study setup with color scheme. Plasma concentration of B) CD4, C) IL-32, D) Reg-3-alpha and E) I-FABP. Two-way repeated measurements ANOVA was used to analyse the effect of acute exercise (EX), bodyweight (BW) and the interaction (EX x BW). Data is shown as mean± SEM with individual participants represented as dots. A was created using BioRender.

## 4.0 Discussion

In this study, we applied an integrative approach combining human plasma analyses and murine experiments to investigate how acute and chronic physical activity, fasting-refeeding, and caloric dilution influence intestinal morphology, transcriptional programs, and systemic markers of epithelial and immune function. Our findings reveal that the intestinal epithelium and immune landscape are highly plastic, adapting in a segment-specific manner to energy demands imposed by exercise and diet. Our data aligns with emerging concepts of the gut as an adaptive organ playing a significant role in systemic energy homeostasis and in the regulation of immunometabolism^47^.

Murine studies allowed us to dissect the tissue-specific mechanisms underlying these systemic changes. Chronic voluntary activity resulted in longer small intestines and increased colonic length, consistent with adaptive remodeling in the small intestine to support higher energy intake and nutrient absorption^44^. Acute exercise, in contrast, elicited transient shortening of the small intestine and rapid upregulation of genes involved in apoptosis, proliferation, and immune regulation^53–56^ *(Bcl2l1, Casp3, Wnt2b, Ki67*) in sedentary mice, while these responses were blunted in chronically active mice. These findings suggest that repeated exercise induces adaptive remodeling, dampening stress and immune activation in response to acute bouts, while maintaining elevated baseline expression of key genes, potentially contributing to intestinal homeostasis and barrier integrity. Similarly, fasting-refeeding experiments revealed that sedentary mice mount robust segment-specific transcriptional responses, particularly in proximal and distal small intestine, whereas chronically active mice showed attenuated or even downregulated responses, notably for *Wnt2b.* The dampening of acute transcriptional activation by chronic exercise may represent a protective adaptation, limiting excessive epithelial turnover and immune signaling in response to nutrient intake, thereby maintaining homeostasis and preventing low-grade inflammation, as has been seen in the post-fasting refeeding^14^.

Chronic caloric dilution further demonstrated the interaction between diet and activity in shaping intestinal adaptation. Caloric dilution feeding induced upregulation of *Ki67, Cyba*, and *Nfil3* in the proximal small intestine, downregulation of *Casp3* in the mid intestine, and downregulation of *Bcl2l1* in the distal segment, reflecting segment-specific compensatory mechanisms in response to reduced energy intake. Chronic exercise primarily influenced the proximal intestine, with trends toward upregulation of *Bcl2l1, Wnt2b,* and *Nfil3*, whereas systemic markers showed tendencies of exercise-induced elevations of CD14 across diets, and I-FABP bouts only in exercised mice on the dilution diet, though high variability among control-fed mice suggests caution in interpretation. Together, these data highlight that dietary challenges and exercise act synergistically and segment-specifically to shape intestinal transcriptional and systemic responses.

Complementary, in humans, acute moderate-intensity exercise led to increased plasma levels of CD14, IL-32, Reg-3-alpha and I-FABP, highlighting rapid systemic responses of both immune and epithelial compartments. Elevated I-FABP reflects small intestinal epithelial damage^48^, which can allow microbial products to cross the weakened barrier and in turn active the innate immune signaling, here seen by elevated CD14 levels^45^, and increased production of IL-32, a cytokine that amplifies the inflammatory response^49,50^. These effects were largely independent of lean or obese conditions, suggesting that the acute intestinal and immune response to exercise is generally conserved albeit we cannot exclude an impact of metabolic status in subjects with more severe obesity and/or with obesity-associated comorbidities^51,52^. Notably, because lean and obese participants exhibited similar absolute VOLpeak (mL/min)^23^, indicating comparable aerobic capacity, the similar magnitude of responses between groups suggests that adiposity alone does not confer intestinal “preconditioning,” and that training status, rather than body composition, is likely the dominant determinant of exercise-induced gut adaptations.

Several limitations of the current study should be considered. All mice in the running group were included regardless of running distance and the present work was not statistically powered, since our primary aim was to generate initial biological insights rather than to evaluate a specific effect-size hypothesis^57^. Moreover, although bulk transcriptomics revealed only modest changes in specific genes after correction for multiple comparisons, we explored the underlying biological mechanisms in all datasets as exploratory analyses. Our analyses were limited to murine bulk transcriptomics and gene expression profiling across multiple cell types within each intestinal segment, which may mask cell-type-specific adaptations. The human analyses included only acute moderate-intensity exercise and, although the plasma biomarkers provide systemic signals, they do not allow segment-specific intestinal analyses, limiting mechanistic insight. Hence, the effect of chronic exercise training in humans remains untested. Finally, we did not examine tissue-level adaptations in females and given the sex-specific differences in response to running^58^, further analyses are required to determine the influence of nutritional status and increased physical activity on the intestinal immune response in females.

Collectively, our findings support a model in which the gut integrates multiple energy signals, including dietary intake, fasting-refeeding cycles, and physical activity, to modulate epithelial turnover, apoptosis, proliferation, and immune signaling in a segment-specific and adaptive manner. Acute exercise and nutrient fluctuations trigger rapid, transient responses in small intestinal length and gene expression, whereas chronic activity and dietary adaptations lead to dampened or reprogrammed responses in parameters related to acute immune-activity, promoting intestinal homeostasis and resilience. These insights have important implications for understanding how lifestyle interventions, including exercise and diet, may improve metabolic health and gut function, and suggest that segment-specific intestinal adaptations are a critical feature of integrated energy homeostasis.

## Author contribution

Conceptualization: C.B-L. and P.S. Methodology: C.B-L., H.E., B.A.H.J, B.K.P and P.S., Formal analysis: C.B-L., J.V., and P.S. Investigation: C.B-L., N.S.N., B.T., C.M., S.V., A.H.P, J.V, and P.S., Writing – original draft: C.B-L. and P.S. Writing – review & editing: C.B-L, N.S.N, B.T., C.M., S.V., A.H.P., H.E., J.V., B.A.H.J., B.K.P., P.S. Visualization: C.B-L. and P.S. Supervision: C.B-L., B.K.P., and P.S. Funding acquisition: B.K.P. and P.S.

## Acknowledgements

Anne Jørgensen, Lene Foged and Ida Holm are acknowledged for their technical assistance. This study was supported by a research grant from the Novo Nordisk Foundation (grant ID 0059436). The Centre for Physical Activity Research (CFAS) is an independent research center at Rigshospitalet and is supported by Trygfonden (grants ID 101390, ID 20045, ID 125132, and ID 177225).

P.S. was awarded a Lundbeck Foundation research grant (grant ID R380-2021-1300). B.T was funded by Research Funds of the University of Basel (grant ID: 3MS1019) and the Swiss National Science Foundation (grant ID: P400PM_191043).

## Materials Availability

This study did not generate new unique materials.

## Data availability

The transcriptomic raw RNA-seq data have been deposited at the Sequence Read Archive Database (https://www.ncbi.nlm.nih.gov/sra/) under BioProject accession PRJNA1210847. https://dataview.ncbi.nlm.nih.gov/object/PRJNA1210847?reviewer=c45r9cbjkrm0oqgb74ub3t0ulv Any additional information required to reanalyze the data reported in this work paper is available from the corresponding author upon request.

## Supplemental information

Document S1. Figures S1-S3

## Supplementary Figures

**Figure S1.** **Volcano plots of differential expressed genes in response to voluntary wheel running**. A) proximal section. B) Middle section. C) Distal section. Genes with p-value<=0.01 and absolute log_2_FC greater than 1, p-values<=0.05 and absolute log2FC greater than 1 and p-value<=0.01 were highlighted in red, pink, and brown, respectively. Non-significant genes were labelled in grey.

**Figure S2.** **Functional protein-protein association networks in each intestinal section**. A) STRING analysis of differential expressed genes from proximal section. B) STRING analysis of differential expressed genes from middle section. C) STRING analysis of differential expressed genes from distal section.

**Figure S3.** Phenotypic analysis previously published, related to. Figure 3. A) Total running distance. B) Total food intake (grams). C) Total food intake (Kcal). D) Body weight changes. E) Lean mass changes. F) Fat mass changes. Effect of calorie dilution diet and of voluntary wheel running was compared by two-way ANOVA with Bonferroni multiple comparison. Bar graphs show mean ± SEM with individual mice represented as dots.

