## Supplementary figures and images for "The intestinal immune response is influenced by nutritional-status and increased physical activity level"

### Suppl figures

Supplementary figure 1

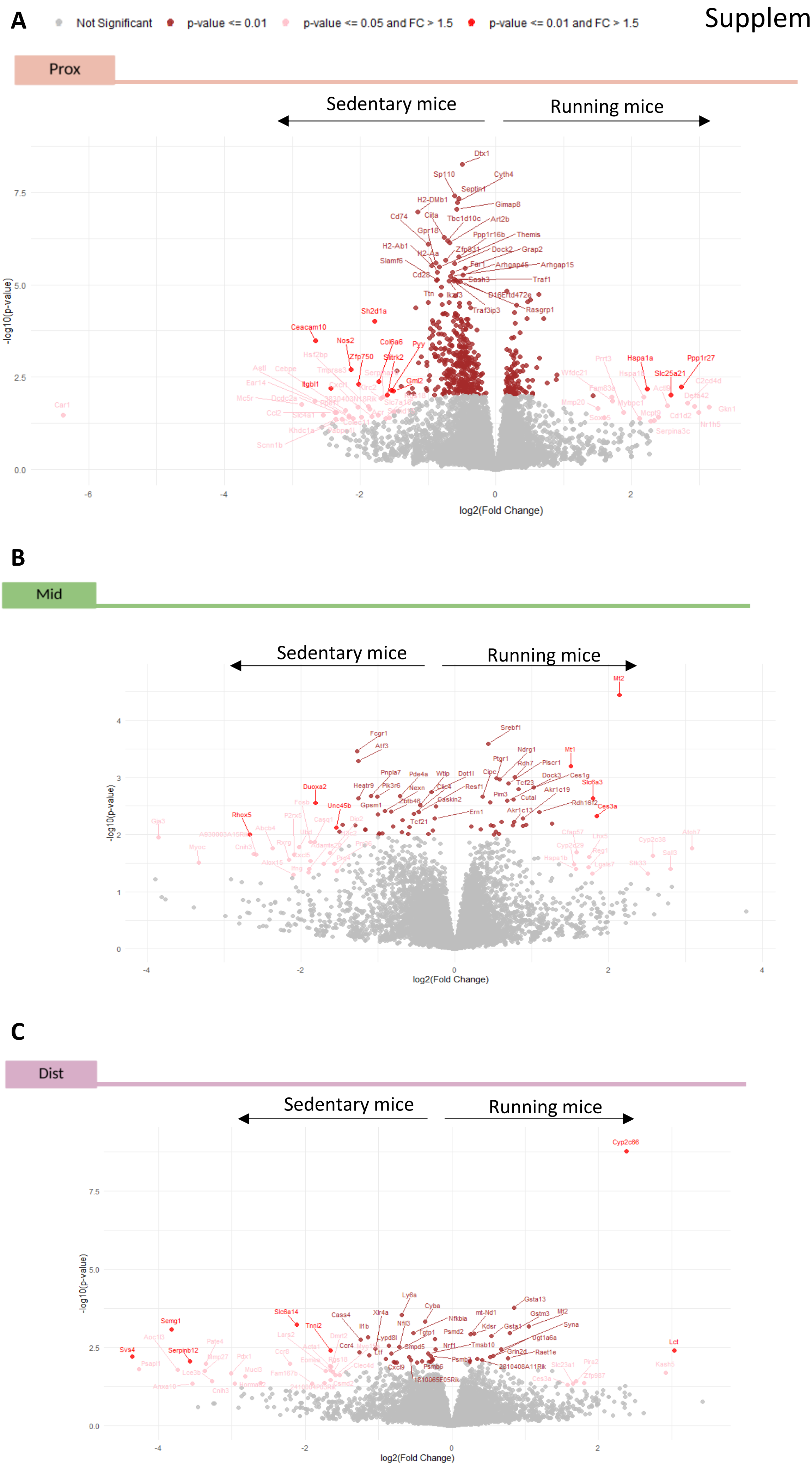

Supplementary figure 2

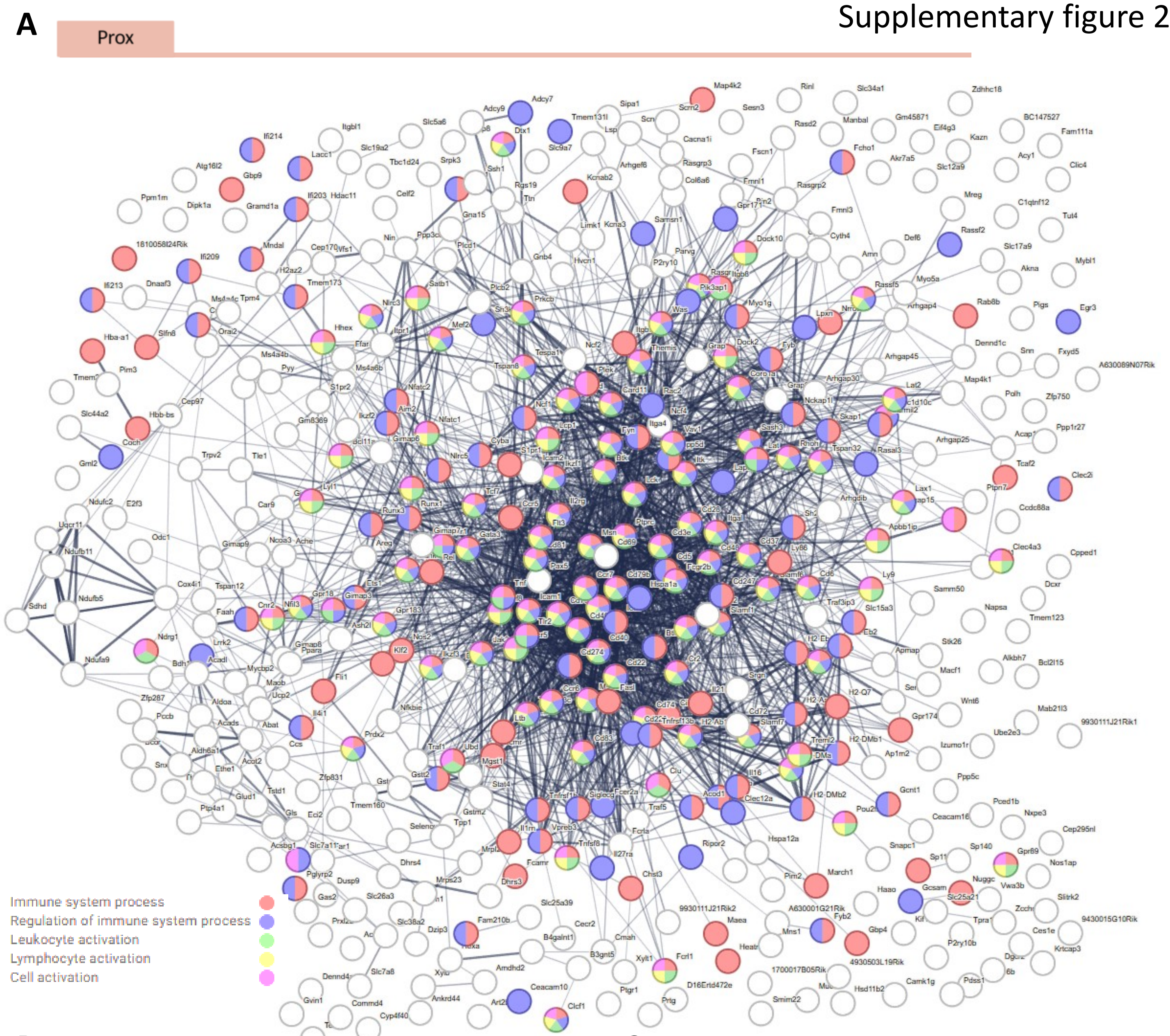


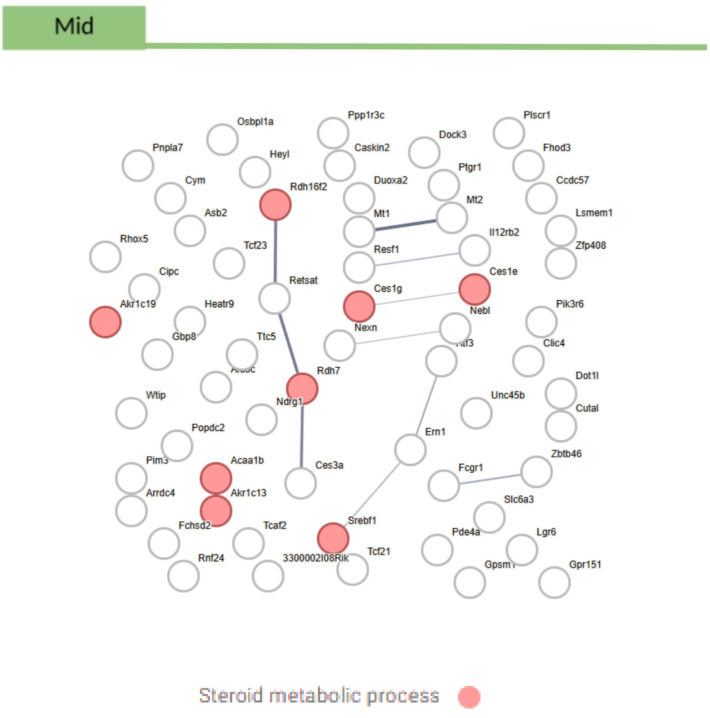


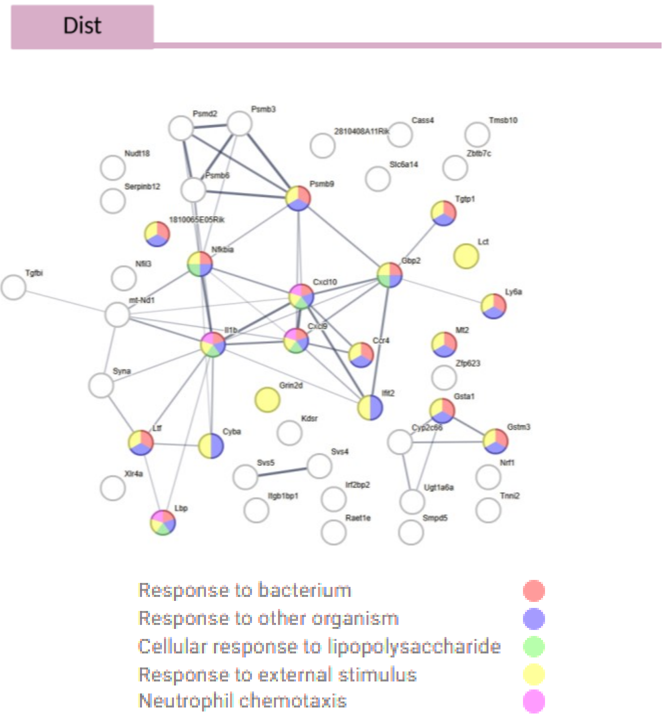

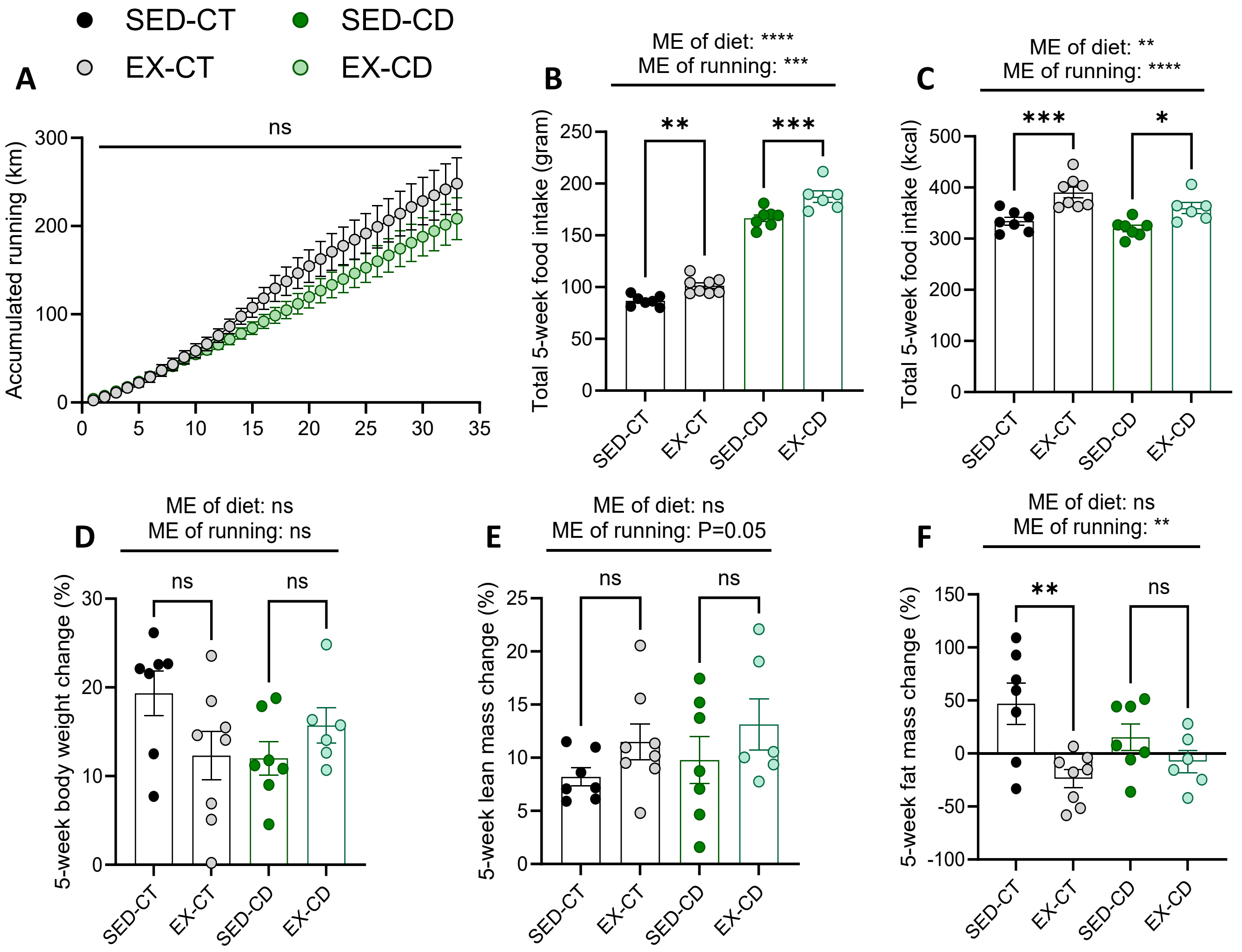
